# A harmonized single-cell RNA-seq atlas of human localized and metastatic prostate cancers and benign tissues

**DOI:** 10.64898/2026.05.18.725966

**Authors:** Hanbyul Cho, Yuping Zhang, Jiayi Zhou, Aniket Daggar, Sarah Kang, Rahul Mannan, Xuhong Cao, Saravana Mohan Dhanasekaran, Arul M. Chinnaiyan

## Abstract

Single-cell RNA sequencing (scRNA-seq) effectively captures the differences in transcriptomic landscape of cell types and cell states between benign and cancer tissues. Pooling publicly available datasets distributed across independent studies enables increased sample representation and cross-study comparisons. Here we present a harmonized scRNA-seq atlas of the human prostate constructed by integrating 17 available studies, comprising 163 samples from 106 donors. The dataset contains benign tissue, primary tumors, and metastatic disease profiles. Raw sequencing FASTQ data files were uniformly reprocessed to minimize technical variability. Study metadata were curated and standardized using a unified schema capturing donor identity, tissue site, disease context, and histologic grade. Post quality control, the integrated dataset contains 754,000 high-quality cells. Harmonized cell type annotations were generated using a pseudobulk correlation framework informed by multiple reference resources. The workflow identified 17 distinct cell types representing epithelial, mesenchymal, and immune compartments of the prostate. The processed expression matrices, standardized metadata, and analysis workflows are publicly available to support reproducible analysis and enable exploration of heterogeneity across prostate disease states.

## Background & Summary

Prostate cancer is one of the most frequently diagnosed malignancies among men worldwide and displays substantial biological and clinical heterogeneity ^1^. Disease progression can span preneoplastic lesions, localized tumors with relatively favorable outcomes, and advanced metastatic disease, including metastatic castration-resistant prostate cancer (mCRPC), which remains largely incurable ^2^. A unified understanding of both the cellular composition and transcriptional programs of cell states underlying prostate tumor progression is therefore critical for improving disease characterization and therapeutic strategies ^3,4^.

Bulk transcriptomic profiling has provided important insights into prostate cancer biology, but these measurements represent averaged signals across heterogeneous cell populations ^5^. As a result, bulk sequencing cannot distinguish transcriptional programs originating from malignant epithelial cells, stromal components, immune populations, or other cell types within the tumor microenvironment. Single-cell RNA sequencing (scRNA-seq) overcomes this limitation by enabling gene expression profiling at cellular resolution and facilitating systematic characterization of cell types and transcriptional states within complex tissues ^5,6^.

In recent years, multiple scRNA-seq studies have profiled human prostate tissues across diverse biological contexts, including normal prostate, benign tissue, benign prostatic hyperplasia (BPH), localized tumors, and advanced disease states ^6–30^. These studies have revealed diverse epithelial, stromal, and immune populations within prostate tissues. However, individual studies typically contain limited numbers of samples and are often processed using different experimental platforms and analytical pipelines, which precludes direct cross-study comparisons.

To address these challenges, integrated single-cell atlases have emerged as valuable community resources that harmonize datasets across studies using standardized computational workflows ^9,31^. Such resources enable cross-study analyses, facilitate benchmarking of computational methods, and provide large-scale references for investigating tissue heterogeneity.

Here we present a harmonized single-cell RNA-sequencing atlas of the human prostate assembled from multiple publicly available scRNA-seq datasets (**Supplementary Table 1**). Raw sequencing FASTQ files were downloaded from publicly available repositories (GEO, SRA, and supplementary repositories associated with each study) and uniformly reprocessed using a standardized computational pipeline. Sample metadata obtained from each study were curated and standardized to facilitate consistent cross-dataset comparisons (**Supplementary Table 2**). The overall workflow for dataset assembly and processing is illustrated in **Figure 1** and includes data collection, metadata harmonization, quality control, cell-type annotation, and dataset integration. The atlas contains over 750,000 high-quality cells and captures 17 distinct annotated cell types spanning the key epithelial, immune, and mesenchymal compartments (**Figure 2**). Cell type proportions, marker gene expression, and cellular subtype distributions within epithelial, immune, and mesenchymal compartments are consistent with known prostate biology (**Figures 3 and 4**).

**Figure 1.**
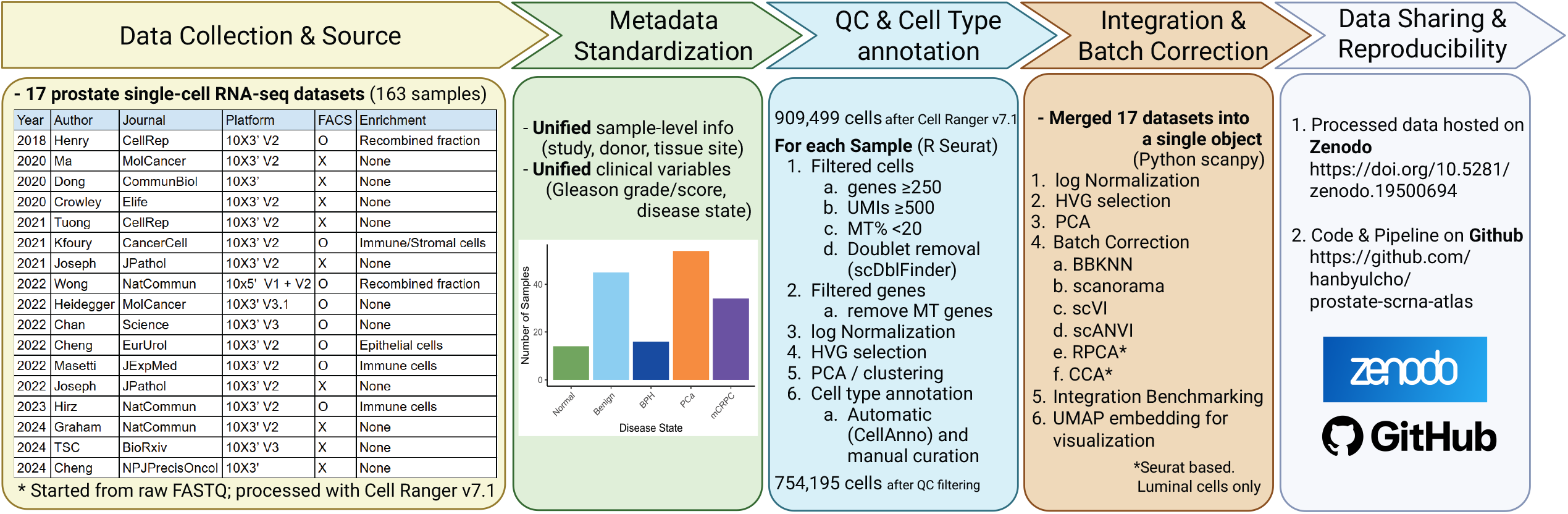
Workflow for assembly and processing of the integrated human prostate single-cell RNA-seq atlas. Publicly available 10x Genomics platform-based prostate scRNA-seq datasets were collected from 17 studies comprising 163 samples from 106 donors (Supplementary Table 1). The samples span benign prostate tissue, primary tumors, and metastatic disease. Raw FASTQ files were uniformly reprocessed using Cell Ranger (v7.1), yielding 909,499 cells prior to quality control. Sample-level metadata were curated and standardized to harmonize variables across studies, including study source, donor identity, tissue site, disease context, and clinical annotations (Supplementary Table 2). For each sample, cells were filtered using thresholds on gene counts (≥250), UMI counts (≥500), mitochondrial transcript fraction (<20%), and doublet detection (scDblFinder). Gene expression matrices were log-normalized and highly variable genes were identified prior to dimensionality reduction and clustering. Cell types were annotated using a pseudobulk correlation framework (CellAnno ^34^ reference expanded with Henry et al. ^7^ prostate profiles, with Tabula Sapiens ^9^ and Human Protein Atlas ^35^ profiles for immune and stromal populations) followed by manual curation; cells labeled as low quality or unknown were excluded, yielding a final dataset of 754,195 high-quality cells. Datasets were subsequently integrated using scANVI, selected based on quantitative benchmarking across six integration methods (Figure 5). Processed data, standardized metadata, and analysis pipelines are publicly shared to facilitate reproducibility and reuse.

**Figure 2.**
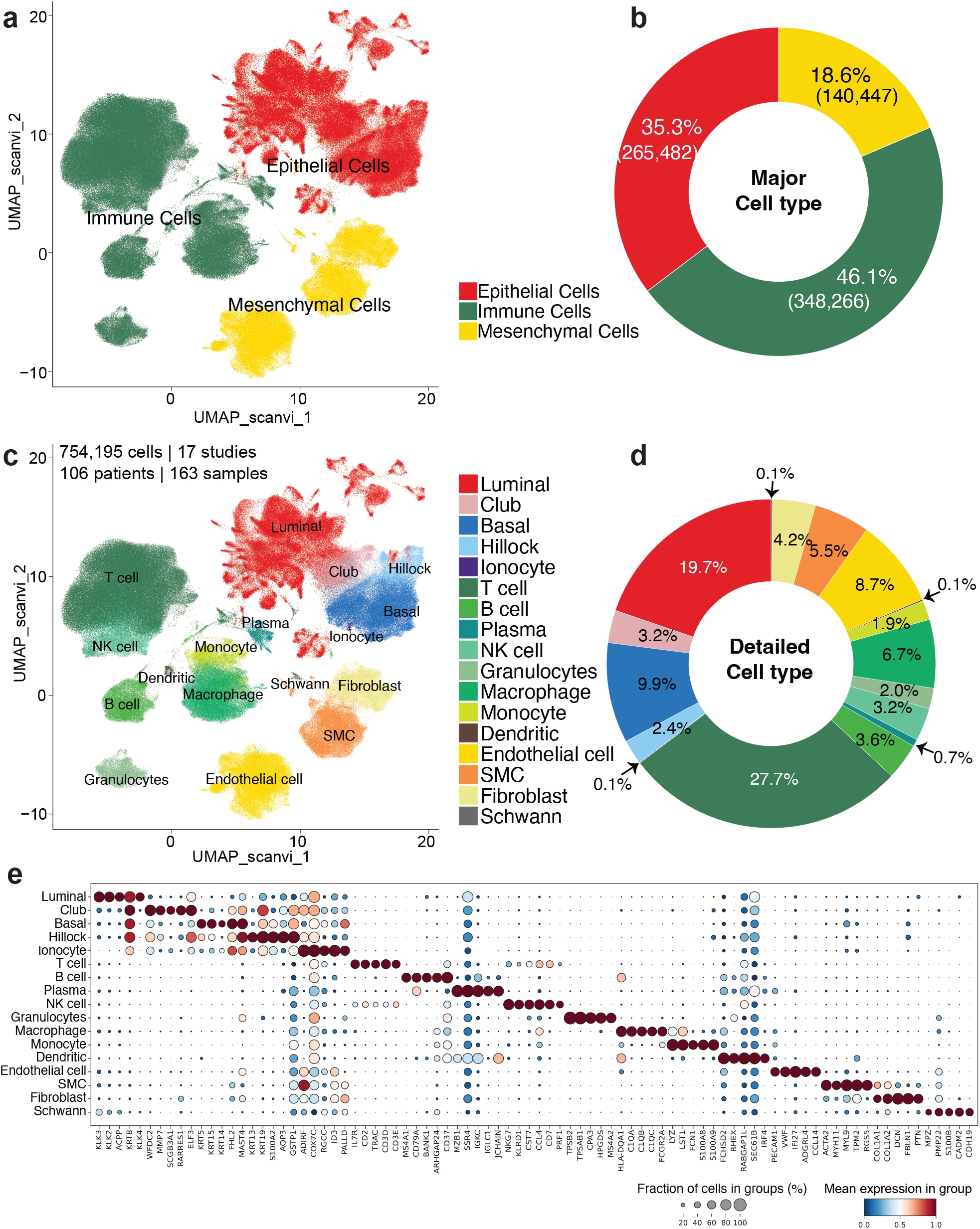
Cell type composition of the integrated human prostate single-cell atlas. **(a)** UMAP embedding of 754,195 cells integrated using scANVI, colored by major cell type compartment: epithelial cells (red), immune cells (green), and mesenchymal cells (yellow). **(b)** Donut chart showing the proportion and absolute cell counts of major cell type compartments: epithelial (35.3%, 265,482 cells), immune (46.1%, 348,266 cells), and mesenchymal (18.6%, 140,447 cells). **(c)** Same UMAP as (a), colored by annotated detailed cell type. Seventeen cell populations are identified spanning epithelial, immune, and mesenchymal compartments. **(d)** Donut chart showing the proportion of individual detailed cell types. T cells constitute the largest single population (27.7%). Among epithelial cells, luminal cells are the most abundant (19.7%), followed by basal cells (9.9%). Among mesenchymal cells, endothelial cells are the most abundant subtype (8.7%). **(e)** Dot plot displaying top marker genes for each annotated cell type. Circle size indicates the fraction of cells expressing the gene; color intensity indicates mean normalized expression.

**Figure 3.**
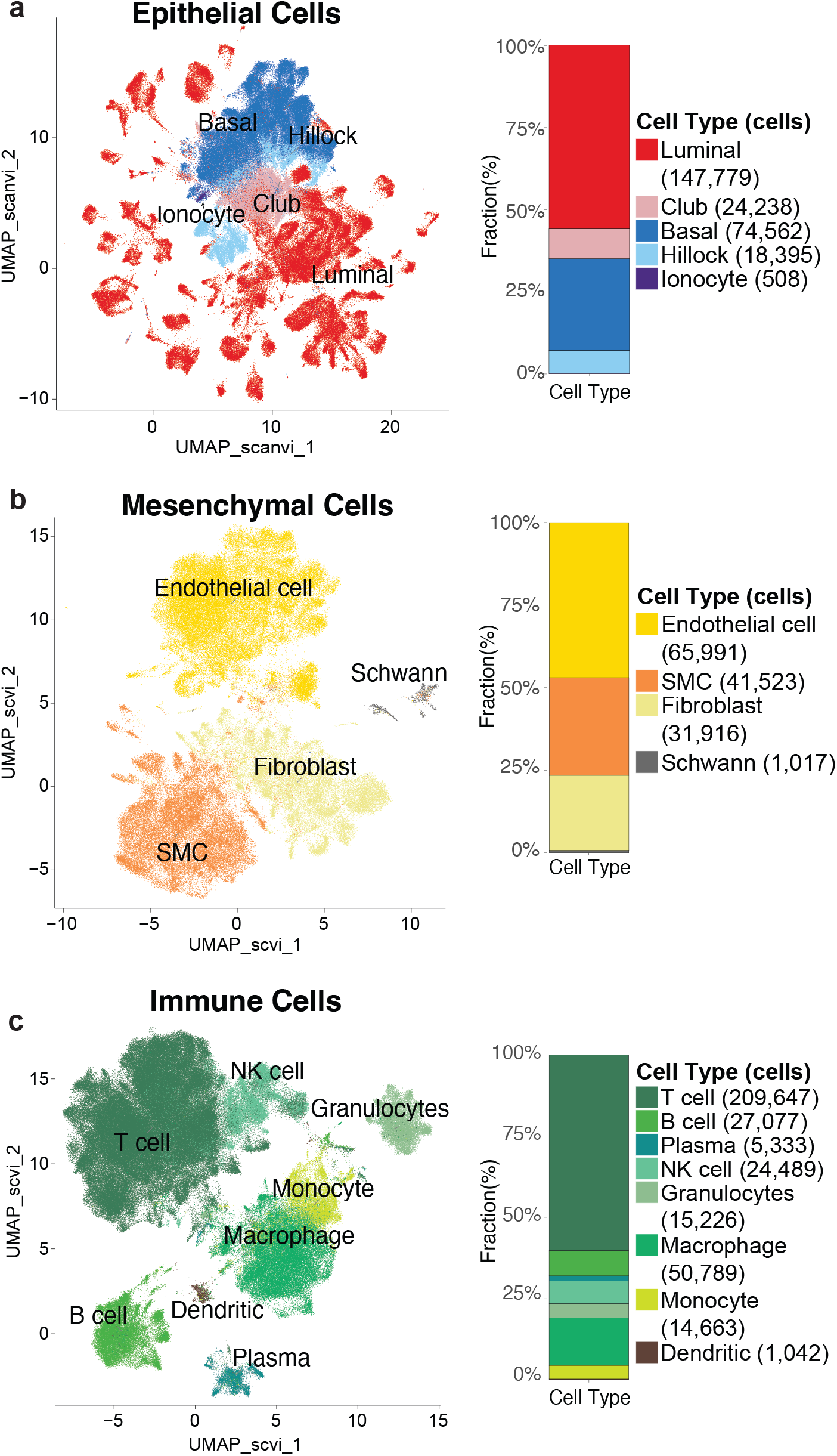
Compartment-level characterization of the integrated atlas. **(a)** UMAP embedding of epithelial cells (scANVI), colored by subtype: luminal, club, basal, hillock, and ionocyte. The bar chart (right) shows the proportional composition and absolute cell counts of epithelial subtypes across the atlas. **(b)** UMAP embedding of mesenchymal cells (scVI), colored by subtype: endothelial cell, SMC, fibroblast, and Schwann. The bar chart (right) shows the proportional composition and absolute cell counts of mesenchymal subtypes. **(c)** UMAP embedding of immune cells (scVI), colored by subtype: T cell, B cell, plasma, NK cell, granulocytes, macrophage, monocyte, and dendritic. Bar chart (right) shows the proportional composition and absolute cell counts of immune subtypes.

**Figure 4.**
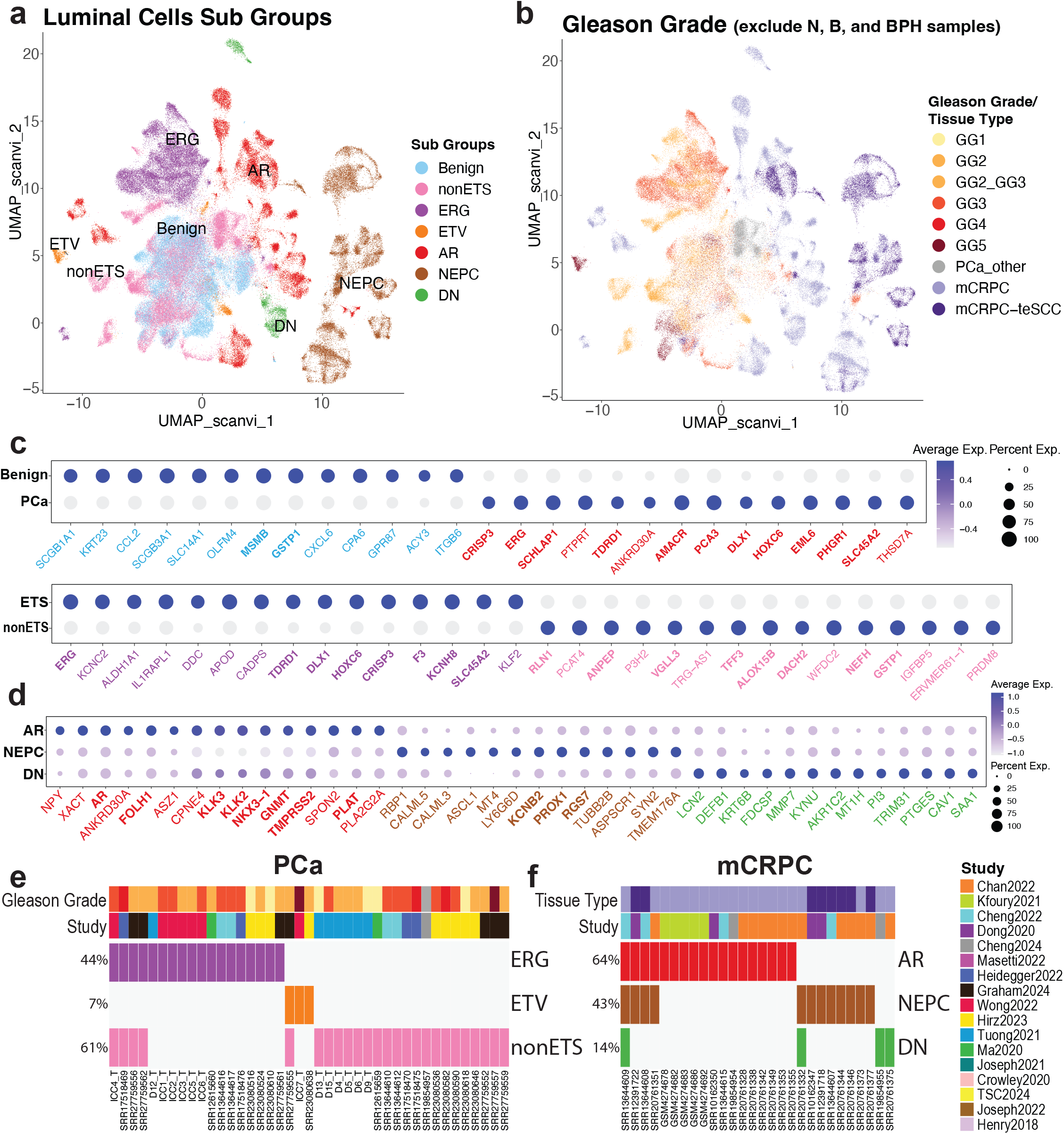
Luminal cell subtype characterization across disease states. **(a)** UMAP embedding of luminal cells colored by transcriptional subgroup: benign-associated (Benign, BPH), ETS fusion-positive primary tumor (ERG, ETV), ETS fusion-negative primary tumor (nonETS), and metastatic castration-resistant prostate cancer subtypes (AR-driven [AR], neuroendocrine [NEPC], and double-negative [DN; AR−/NE−]). **(b)** Same UMAP as (a), colored by Gleason grade for primary tumor samples and tissue type for mCRPC samples. Cells from normal (N), benign (B), and BPH samples are excluded. **(c)** Dot plots showing marker gene expression distinguishing benign versus primary prostate cancer (PCa) luminal cells (top) and ETS-positive versus ETS-negative (nonETS) tumor cells (bottom). Canonical marker gene names are bolded and colored by their corresponding transcriptional subgroup, matching the color scheme in (a). Circle size indicates the percentage of cells expressing the gene; color intensity indicates scaled average expression. **(d)** Dot plot showing marker genes for mCRPC subtypes (AR, NEPC, DN; AR−/NE−), with canonical marker gene names bolded and colored by subgroups as in (c). **(e)** Sample-level stacked bar plots showing proportional composition of ETS subgroups (ERG, ETV, nonETS) across primary prostate cancer samples. Metadata tracks indicate Gleason grade and study of origin. **(f)** Same as (e) for mCRPC samples showing AR, NEPC, and DN subgroup composition with tissue type and study metadata tracks.

Luminal epithelial populations were further resolved into transcriptionally distinct subgroups, including benign-associated luminal cells and tumor luminal cells spanning ETS fusion-positive (ERG, ETV), ETS fusion-negative (nonETS), and mCRPC subtypes (AR-driven, neuroendocrine prostate cancer (NEPC), and double-negative), characterized by subgroup-specific marker gene expression and inferred copy number variation (**Figure 4**). The resulting integrated atlas captures cell types from epithelial, mesenchymal, and immune cell compartments found in human benign and malignant prostate tissues. Processed data (batch corrected, **Figure 5**), standardized metadata, and analysis pipelines are publicly deposited at Zenodo (https://doi.org/10.5281/zenodo.19500694) and GitHub (https://github.com/hanbyulcho/prostate-scrna-atlas) to support transparency, reproducibility, and reuse. Utilization of this large atlas will increase statistical power for detecting rare cell populations and enable systematic comparisons across disease states and experimental platforms. This dataset will be a valuable resource for the research community to explore disease mechanisms, prostate cellular heterogeneity, and transcriptional regulation, and to support future studies of prostate biology and disease. This dataset will also facilitate benchmarking of computational methods and incorporation into pan-cancer studies.

**Figure 5.**
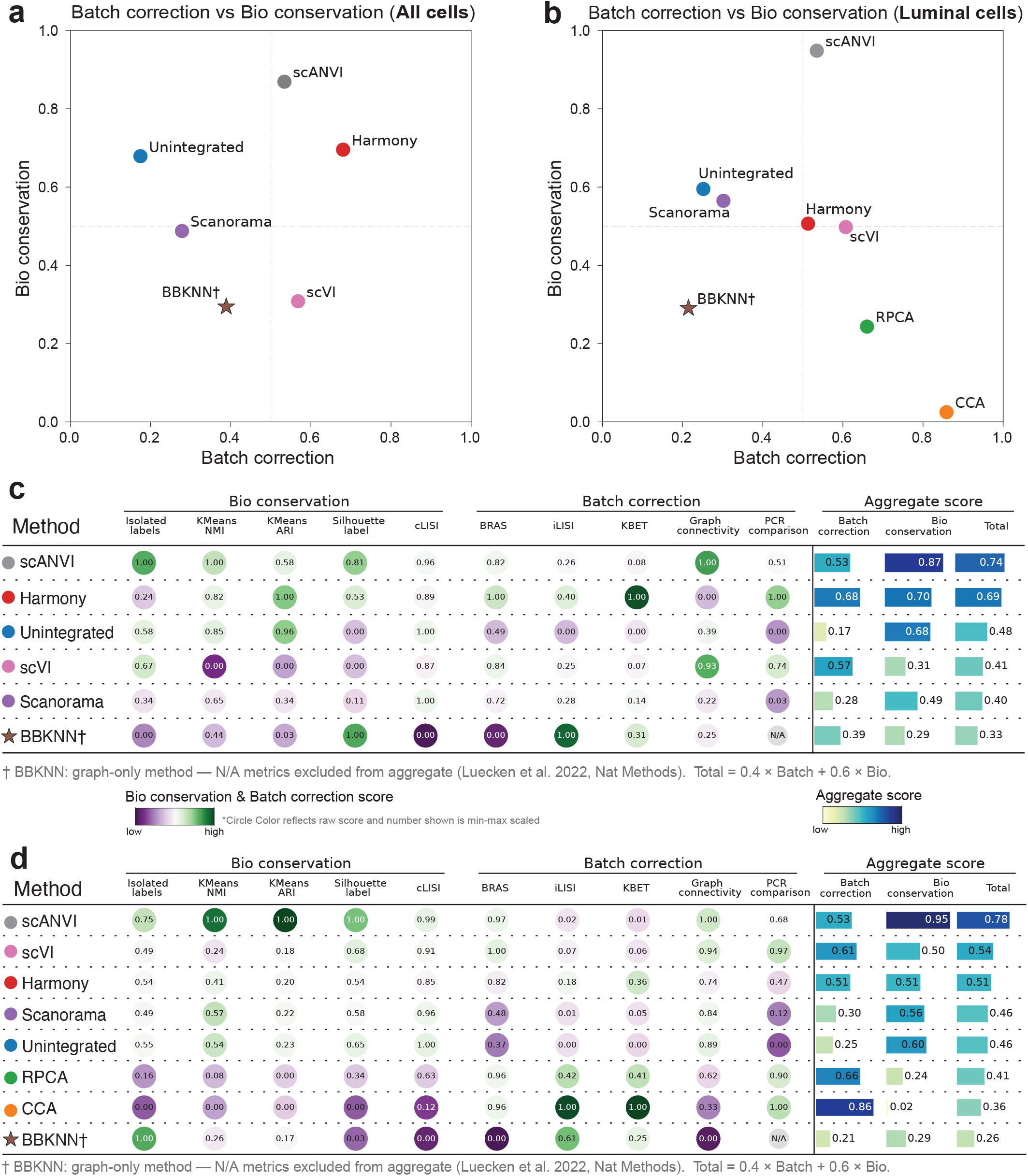
Benchmarking of data integration methods for the human prostate single-cell atlas. **(a)** Scatter plot comparing min-max scaled batch correction (x-axis) and biological conservation (y-axis) aggregate scores for six integration methods applied to all cell types (2,000 highly variable genes, 50 principal components). Each point represents one method. Aggregate sub-scores are means of five per-column min-max scaled metrics each (0 = worst, 1 = best method per metric). Total score = 0.4 × Batch correction + 0.6 × Bio conservation. Dashed lines indicate a score of 0.5. Methods in the upper-right quadrant achieve both effective batch integration and preservation of biological structure. **(b)** Same as (a) for luminal cells only (3,000 highly variable genes, 30 principal components), additionally including Seurat-based integration methods CCA and RPCA. **(c)** Detailed scib benchmark scores for all cell types. Circles display individual metric scores for biological conservation (Isolated labels, KMeans NMI, KMeans ARI, Silhouette label, cLISI) and batch correction (BRAS, iLISI, KBET, Graph connectivity, PCR comparison). Circle color reflects the raw metric score per column (purple = low, green = high). Numbers shown within circles are min-max scaled scores (0 = worst, 1 = best method in that column). Aggregate scores (right) are displayed as horizontal bars (light = low, dark blue = high). Methods are ranked by descending total aggregate score. †BBKNN is a graph-only method; PCR comparison requires a corrected high-dimensional embedding and is therefore not applicable; it is excluded from the aggregate score. **(d)** Same as (c) for luminal cells only, additionally including CCA and RPCA.

## Methods

### Data collection

Our unified workflow schematic lists the details of this process (**Figure 1**). Briefly, publicly available human prostate single-cell RNA sequencing (scRNA-seq) datasets were collected from the Gene Expression Omnibus (GEO), Sequence Read Archive (SRA), and supplementary repositories associated with published studies. Two datasets were obtained from controlled-access repositories: Tuong et al. from the European Genome-phenome Archive (EGAS00001005787) and Graham et al. from dbGaP (phs003480.v1.p1). Access to these datasets was granted through approved data access requests in accordance with NIH Genomic Data Sharing policies and the data use agreements specified by the original data providers. All datasets were reused under their respective data sharing terms. As inclusion criteria, we only considered 10x Genomics datasets that contained both raw FASTQ files and accompanying metadata describing tissue origin and disease state. In total, 24 prostate scRNA-seq datasets were initially identified and evaluated for inclusion (**Supplementary Table 1**). Seven datasets were excluded for the following reasons: (1) raw FASTQ files could not be obtained despite more than one year of negotiation with the corresponding authors ^24,25^; (2) data were inaccessible due to database availability and download restrictions ^26–28^; or (3) sequencing was performed on non-10x Genomics platforms (e.g., GEXSCOPE, SeqWell, SMRT-seq), which were incompatible with the harmonized Cell Ranger v7.1 preprocessing pipeline ^6,29,30^. The remaining 17 datasets, comprising 172 samples from 111 donors, were included for processing. Of these, 9 samples were subsequently excluded during quality review for reasons including low feature counts or absence from the original paper’s supplementary metadata (**Supplementary Tables 1, 2**), yielding a final dataset of 163 samples from 106 donors. All datasets were generated using 10x Genomics single-cell platforms across multiple chemistry versions (v2, v3, and v3.1). For consistency across studies, all datasets were reprocessed starting from the raw FASTQ files using a unified computational workflow. Study-level information including publication source, sequencing platform, and sample counts is summarized in **Supplementary Table 1**.

### Raw sequencing processing

Raw sequencing reads were processed using Cell Ranger (v7.1, 10x Genomics) with default parameters. Reads were aligned to the human reference genome (GRCh38) and gene-cell count matrices were generated for each sample. This step produced an initial dataset comprising approximately 909,499 cells across all samples prior to quality control filtering.

### Quality control

Quality control was performed using Seurat (v5.1) ^32^ in R. Cells were filtered according to standard single-cell quality metrics to remove low-quality droplets and potential artifacts. Cells were retained if they satisfied the following criteria: ≥250 detected genes per cell, ≥500 total UMI counts, <20% mitochondrial transcript content. Genes detected in very few cells were removed from downstream analyses. Potential doublets were identified and removed using scDblFinder (v1.16.0) ^33^, which detects multiplet profiles based on simulated doublet distributions. After quality filtering and doublet removal, 767,524 singlet cells were retained. Following cell type annotation, cells labeled as low quality or unknown were excluded, yielding a final dataset of 754,195 high-quality cells (**Supplementary Figure 1, Supplementary Table 2**).

### Metadata curation and standardization

Metadata from individual studies were manually curated and standardized to create a unified metadata schema across datasets. Sample-level metadata fields included: study identifier, donor identifier, tissue origin, disease state, and clinical annotations (e.g., Gleason score or grade group when available) (**Supplementary Table 2**). Because cell type annotations differed across studies, previously published annotations were harmonized into a unified ontology informed by established prostate cell type definitions ^7,34,35^; cell labels from other datasets were mapped to this standardized annotation schema.

### Cell type annotation

Cell type annotation was performed using a prostate-specific, cluster-level pseudobulk annotation framework implemented with Seurat (v5.1) ^32^ and SingleR (v2.4.1) ^36^ in R. A prostate-focused reference was first constructed from the CellAnno transcriptomic reference resource ^34^ (https://github.com/yupingz/CellAnno), which includes prostate epithelial, stromal, and immune cell populations, with additional reference profiles from the Tabula Sapiens ^9^ and Human Protein Atlas ^35^ incorporated for selected immune and stromal populations. This reference was then expanded using pseudobulk profiles derived from the normal human prostate dataset ^7^, generating an updated prostate-specific reference used for final annotation. Within each sample, cells were clustered and cluster-level pseudobulk expression profiles were generated by aggregating counts within clusters. These profiles were normalized and compared against the reference using SingleR ^36^, and predicted labels were transferred back to individual cells based on cluster membership. A cluster-level pseudobulk strategy was used to improve annotation stability and robustness in this large integrated dataset. Epithelial subsets were subsequently re-clustered and re-annotated to improve resolution of luminal, basal, club, and hillock populations, and final labels were refined using canonical marker genes and manual review. Final annotated cell populations and their marker gene signatures are summarized in **Figure 2a-e**, and compartment-level distributions across samples are shown in **Figure 3a-c**. A further detailed characterization of the luminal cell compartment based on marker expression patterns is presented in **Figure 4a-f**.

### Data integration and batch correction

Following quality control and annotation using Seurat (v5.1) ^32^ in R (v4.3.1), the filtered dataset was exported and processed using a Python-based workflow (Python v3.10) implemented with Scanpy (v1.10.0) ^37^ and anndata (v0.10.5) ^38^. For each study, sample-level 10x Genomics gene-barcode matrices were loaded and merged into a study-level dataset. Study-level datasets were then concatenated into a single AnnData object using Paper_ID as the batch identifier. Cells were restricted to those present in the curated annotation table; cells labeled as low quality or unknown were excluded. Mitochondrial genes were removed prior to downstream analyses. Raw counts were stored as a dedicated layer, and expression matrices were total-count normalized (target sum 10,000) and log-transformed prior to dimensionality reduction. Highly variable genes (HVGs) were selected in a batch-aware manner using Paper_ID as the batch key, using the Seurat (v5.1) ^32^ HVG selection method applied to raw counts.

For benchmarking across all cell types, 2,000 HVGs and 50 principal components (PCs) were used. For the luminal cell subset, 3,000 HVGs and 30 PCs were used to better resolve finer transcriptional structure within this compartment. The following integration methods were evaluated both for all cells and luminal cells (**Figure 5a-d**):

⊠ **Unintegrated (PCA baseline):** Standard PCA on scaled HVG expression without batch correction, serving as an unintegrated reference.
⊠ **Harmony (v0.0.9)^39^** Iterative cluster-based batch correction applied to the PCA embedding using harmonypy.
⊠ **Scanorama (v1.7.4):^40^** Panoramic stitching of datasets in PCA space, producing a shared low-dimensional embedding.
⊠ **BBKNN (Batch Balanced k-Nearest Neighbors; v1.6.0) ^41^:** Graph-based batch correction that constructs a balanced nearest-neighbor graph across batches without producing a corrected low-dimensional embedding; UMAP coordinates derived from the BBKNN graph were used as a proxy embedding for benchmarking purposes.
⊠ **scVI (scvi-tools v1.1.2) ^42^:** A deep generative model trained on raw counts (n_latent = 30, n_layers = 2, max_epochs = 400 with early stopping) using Paper_ID as the batch covariate.
⊠ **scANVI (scvi-tools v1.1.2) ^42^:** A semi-supervised extension of scVI incorporating cell type label information (n_latent = 30, max_epochs = 20, n_samples_per_label = 100), initialized from the trained scVI model.

For the luminal cell analysis, two additional Seurat-based integration methods were included: CCA (Canonical Correlation Analysis) and RPCA (Reciprocal PCA), whose corrected embeddings were exported from R and incorporated into the benchmarking workflow. UMAP embeddings were computed for each integration output (n_neighbors = 30, kNN graph k=15) to enable visual assessment of batch mixing and cell type separation.

### Integration benchmarking (scib-metrics; v0.4.1)

Integration performance was quantitatively evaluated using the scib-metrics benchmarking framework ^43^, which simultaneously assesses batch effect removal and preservation of biological structure. Benchmarking was conducted independently for all cell types and for luminal cells. Five biological conservation metrics were computed: Isolated label score, KMeans NMI, KMeans ARI, Silhouette label score, and cLISI. Five batch correction metrics were computed: BRAS, iLISI, kBET, Graph connectivity, and PCR comparison. PCR comparison requires a corrected high-dimensional embedding and is not applicable to BBKNN, and was therefore excluded from its aggregate score. An aggregate total score was calculated as a weighted sum (Total = 0.4 × Batch correction + 0.6 × Bio conservation) following the weighting scheme of Luecken et al. Method comparison was based on per-column min-max scaled scores, which rescale each metric independently from 0 (worst-performing method) to 1 (best-performing method), following the evaluation approach of Luecken et al. Complete raw and min-max scaled metric scores for all cell types and luminal cells are provided in **Supplementary Tables 4-7**. Based on benchmarking results (see Technical Validation), scANVI was selected as the primary integration method for the final atlas.

## Data Record

The deposit contains the following files, publicly available at https://doi.org/10.5281/zenodo.19500694:

1. **All-cells AnnData and Seurat objects**. A pre-integration object (all_cells_raw.h5ad; 754,195 cells after QC and annotation filtering, with raw counts stored as a dedicated layer and log-normalized expression in X) and a multi-method integrated object (all_cells_integrated.h5ad), which stores corrected embeddings and UMAP coordinates from all six benchmarked integration methods (Unintegrated PCA, Harmony, Scanorama, BBKNN, scVI, and scANVI) alongside raw counts and normalized expression. The corresponding Seurat RDS (all_cells_integrated.rds) is also provided.
2. **Luminal-cell AnnData and Seurat objects**. A pre-Python-integration object (luminal_cells_raw.h5ad), which contains CCA, RPCA, and Harmony corrected embeddings from Seurat v5 batch correction alongside raw counts and log-normalized expression, and a multi-method integrated object (luminal_cells_integrated.h5ad), which stores corrected embeddings and UMAP coordinates from all eight benchmarked methods (Unintegrated PCA, CCA, RPCA, Harmony, Scanorama, BBKNN, scVI, and scANVI) alongside raw counts and normalized expression. The corresponding Seurat RDS (luminal_cells_integrated.rds) is also provided.
3. **Cell-level metadata**. Per-cell obs metadata for the all-cells and luminal objects (all_cells_metadata.csv and luminal_cells_metadata.csv, respectively), including cell type labels, batch information, QC metrics, and clinical annotations. The master cell annotation table (all_cells_prefilter_annotation.csv) contains all 767,524 singlet cells prior to annotation-based filtering, including cells subsequently excluded. Column definitions for all metadata fields are provided in Supplementary Table 2.
4. **Quality control metrics**. Per-sample QC summary (QC_summary_sample_level.csv, Supplementary Table 3) and per-cell pre-filter QC metrics (QC_metrics_prefilter.csv).
5. **-**-max scaled scib-metrics scores for all-cells integration (Supplementary Tables 4–5) and luminal-cell integration (Supplementary Tables 6–7).

## Technical Validation

Post dataset assembly and processing, quality was assessed at multiple levels including sequence data quality (total UMIs, mitochondrial read level, doublet detection among others), cell type annotation, cross-study integration, luminal subgroup characterization, and integration benchmarking as detailed in the methods section (**Figure 1**).

Following the QC filtering steps described in the Methods, the initial dataset of 909,499 cells was reduced to 767,524 singlet cells after doublet removal, and to a final 754,195 high-quality cells after annotation-based filtering (**Supplementary Figure 1, Supplementary Table 3**).

### Cell type composition and annotation

Following scANVI-based integration, UMAP visualization confirmed effective mixing of cells from all 17 studies while preserving separation of biologically distinct cell populations (**Figure 2a, c**; **Supplementary Figure 2a**). The final atlas of 754,195 cells comprises 17 annotated cell populations spanning three major compartments: epithelial (35.3%), immune (46.1%), and mesenchymal (18.6%) (**Figure 2b, d**). T cells constitute the largest single population (27.7%). Within the epithelial compartment, luminal cells are the most abundant (19.7%), followed by basal (9.9%), club (3.2%), hillock (2.4%), and ionocyte (0.1%) cells (**Figure 2d**). The luminal compartment encompasses both benign-associated luminal cells and transcriptionally distinct tumor luminal subpopulations, including ETS fusion-positive (ERG, ETV), ETS fusion-negative (nonETS), and mCRPC subtypes (AR-driven, NEPC, and double-negative [DN; AR−/NE−]), as described in detail below (**Figure 4**). All 17 studies contribute cells distributed across shared regions of the UMAP without evidence of study-specific clustering (**Figure 2c; Supplementary Figure 2a, b**).

Cell type annotations were validated against established marker genes for pan-tissue stromal and immune compartments and prostate-specific epithelial populations (**Figure 2e**). Epithelial populations were distinguished by canonical mRNA markers including *KLK3* and *KLK2* (luminal), *SCGB1A1* and *SCGB3A1* (club), *KRT5* and *TP63* (basal), *KRT13* (hillock), and *FOXI1* (ionocyte). Immune lymphoid populations were identified by *CD3D* and *CD3E* (T cells), *CD79A* and *MS4A1* (B cells), *IGHG1* (plasma), and *NKG7* (NK cells). Myeloid cell types were annotated using *S100A8* and *S100A9* (granulocytes and monocytes), *C1QA* and *C1QB* (macrophages), and *HLA-DRA* (dendritic cells). Mesenchymal populations were annotated using *PECAM1* and *VWF* (endothelial cells), *ACTA2* and *MYH11* (smooth muscle cells), *DCN* and *LUM* (fibroblasts), and *S100B* and *PLP1* (Schwann cells).

At the sample level, all three major compartments — epithelial, immune, and mesenchymal — were captured across tissue types and disease states, with proportional variation reflecting both biological differences and study-level cell enrichment strategies (**Supplementary Figure 3a**). Notably, several studies applied targeted enrichment for specific cell fractions (e.g., immune or epithelial cells), which contributes to the observed inter-study variation in compartment composition. Within the epithelial compartment, the proportion of luminal cells was notably higher in mCRPC samples compared to benign and primary tumor samples, consistent with the expansion of tumor luminal populations in metastatic disease (**Supplementary Figure 3b**). Compartment-specific re-embedding confirmed clear internal structure within each compartment: the epithelial embedding revealed distinct luminal, basal, club, hillock, and ionocyte populations; the mesenchymal embedding showed separation of endothelial cells, smooth muscle cells, fibroblasts, and Schwann cells; and the immune embedding distinguished T cells, B cells, plasma cells, NK cells, granulocytes, macrophages, monocytes, and dendritic cells (**Figure 3a–c**).

### Luminal subgroup characterization

Within the luminal epithelial compartment, cells from benign tissue and primary and metastatic tumors co-exist but occupy transcriptionally distinct regions of the embedding. Since the reported scRNA-seq studies seldom provide molecular driver status for tumor samples, we applied an inferential approach to assign putative driver labels among tumor luminal cells based on tumor biomarker expression profiles. In prostate cancer, ETS family gene fusions are known to be mutually exclusive oncogenic drivers that result in outlier mRNA expression of the fusion partner gene. By systematically examining ETS family gene expression in tumor epithelial cells, a putative driver classification can be assigned. Using this logical approach, luminal cells were further resolved into transcriptionally distinct subgroups corresponding to benign tissue (Benign, BPH), ETS fusion-positive primary tumors (ERG, ETV), ETS fusion-negative primary tumors (nonETS), and mCRPC subtypes, including AR-driven, neuroendocrine (NEPC), and double-negative (DN; AR−/NE−) populations (**Figure 4a**). UMAP visualization colored by Gleason grade confirmed grade-associated spatial structure within the tumor compartment, with higher-grade tumors occupying distinct embedding regions (**Figure 4b**). Marker gene dot plots confirmed selective expression of canonical subgroup-defining genes for both primary prostate cancer (PCa) (**Figure 4c**) and mCRPC (**Figure 4d**) luminal subpopulations, including *ERG* and *ETV1/4* (ETS subgroups), *AR* and *KLK2/3* (AR-driven mCRPC), and neuroendocrine markers *KCNB2* and *PROX1* (NEPC). At the sample level, subgroup proportions were consistent with known disease biology: among primary prostate cancer samples, ERG fusion-positive tumors comprised 44% and nonETS tumors the majority at 61%; among mCRPC samples, AR-driven samples predominated (64%), followed by NEPC (43%) and DN (14%) subgroups (**Figure 4e, f**). Pseudobulk copy number variation scores increased progressively with disease grade, with mCRPC samples showing the highest copy number burden (**Supplementary Figure 4c**).

Feature plots of AR transcriptional activity score (AR_TCGA_score), neuroendocrine score (NE_Beltran_score), and key marker genes (AR and ERG) confirmed spatial concordance between subgroup identity and molecular marker expression on the luminal UMAP visualization (**Supplementary Figure 4d**).

### Integration benchmarking

Cross-study integration, embedding generation, and benchmarking were performed in Python using Scanpy/anndata with additional integration tools including Scanorama, BBKNN, and scvi-tools (scVI and scANVI). Across all cell types, scANVI achieved the highest total aggregate score (0.74), followed by Harmony (0.69) (**Figure 5a, c; Supplementary Tables 4, 5**). For luminal cells, scANVI again achieved the highest total aggregate score (0.78), followed by scVI (0.54) (**Figure 5b, d; Supplementary Tables 6, 7**). These results reflect scANVI’s advantage in both batch correction and preservation of biological structure through label-guided integration.

## Usage Notes

Detailed usage notes, including instructions for loading and processing the deposited data files, are available in the project GitHub repository (https://github.com/hanbyulcho/prostate-scrna-atlas).

## Code Availability

All analysis scripts are available at https://github.com/hanbyulcho/prostate-scrna-atlas.

## Data Availability

All processed datasets are available at https://doi.org/10.5281/zenodo.19500694.

## Acknowledgements

We wish to thank the authors of the individual studies utilized here for making their data publicly available. We thank Dr. Stephanie Miner for editing and manuscript submission. We thank Dr. Chandan Kumar-Sinha for the discussion.

## Figure Legends

**Supplementary Figure 1. Quality control metrics and cell counts across processing stages. (a)** Violin plots showing per-cell quality control metrics across 17 studies prior to filtering: UMI counts (nCount_RNA, log10 scale), detected gene counts (nFeature_RNA, log10 scale), and mitochondrial transcript fraction (Percent.mt). Red dashed lines indicate filtering thresholds applied per sample (UMIs ≥500, genes ≥250, MT% <20). Pink shading indicates the excluded region below (UMI, gene) or above (MT%) each threshold. **(b)** Boxplot of per-sample doublet rates (%) estimated by scDblFinder across all 17 studies. Each point represents one sample. **(c)** Bar plot showing the number of cells at each processing stage for each of the 17 studies. Bars represent cells at four stages: before QC filtering (orange), after QC filtering (yellow), after doublet removal (grey), and cells retained in the final dataset after annotation-based filtering (dark green). Total cells: 909,499 before QC; 847,090 after QC filtering; 767,524 after doublet removal; 754,195 in the final dataset. Sample-level counts are provided in Supplementary Table 2.

**Supplementary Figure 2. Study-level cell distribution across the integrated atlas. (a)** UMAP embedding colored by study of origin, illustrating mixing of cells from all 17 datasets across the embedding. **(b)** Donut chart showing the proportion of cells contributed by each study. Hirz2023 contributes the largest fraction (26.1%), followed by Joseph2021 (13.9%) and Graham2024 (9.2%).

**Supplementary Figure 3. Sample-level cell type composition across all 163 samples. (a)** Sample-level stacked bar plot showing the proportional composition of broad cell compartments (epithelial, immune, and mesenchymal) across all 163 samples. Metadata tracks above indicate tissue type, Gleason grade, study of origin, recombined fraction status, and total cell count per sample. Samples are ordered by tissue type and disease state. **(b)** Same layout as (a), showing proportional composition of epithelial cell subtypes (luminal, club, basal, hillock, ionocyte) per sample.

**Supplementary Figure 4. Additional characterization of luminal cell subgroups. (a)** UMAP of luminal cells colored by study of origin. **(b)** Same UMAP colored by tissue of origin, illustrating anatomical source diversity within the mCRPC compartment. **(c)** Boxplot of pseudobulk-level CNV scores across disease/grade categories (N, B, BPH, GG1–GG5, PCa_other, mCRPC, mCRPC-teSCC). Each point represents one pseudobulk unit; a single sample may contribute multiple points reflecting transcriptionally distinct clusters. Points are colored by study. mCRPC samples show the highest CNV burden. **(d)** Feature plots on the luminal UMAP showing AR transcriptional activity score (AR_TCGA_score), neuroendocrine score (NE_Beltran_score), and individual marker gene expression (ERG and AR). Color scale: light grey (low) to dark red (high).

## Supplementary Table Legends

**Supplementary Table 1. Dataset collection summary**. Two-tab Excel workbook summarizing the prostate single-cell RNA-seq datasets evaluated for inclusion in this atlas. **Worksheet 1** — PaperInfo_AllCollection: Records for all 24 datasets initially identified and evaluated. Includes study identifier, first author, journal, publication year, GSE/GEO accession number, DOI, sequencing platform, and reason for exclusion where applicable. Seven datasets were excluded from the final analysis for the following reasons: (1) raw FASTQ files could not be obtained despite more than one year of negotiation with the corresponding authors ^24,25^; (2) data were inaccessible due to database availability and download restrictions ^25–27^; or (3) sequencing was performed on non-10x Genomics platforms (e.g., GEXSCOPE, SeqWell, SMRT-seq), which were incompatible with the harmonized Cell Ranger v7.1 preprocessing pipeline ^6,29,30^. **Worksheet 2** — PaperInfo_CurrentCollection: Detailed summary of the 17 datasets included in the final atlas. For each study, it lists the number of cells and samples obtained from the original publication alongside the number retained in this analysis. Sample counts are presented as used(collected) — for example, 163(172) indicates that 163 samples were used in the analysis out of 172 originally collected across all 17 papers, with 9 samples excluded for reasons detailed in Supplementary Table 2. Numbers of samples per disease state (Normal, Benign, BPH, PCa, mCRPC) are also provided.

**Supplementary Table 2. Sample-level metadata**. Comprehensive metadata for all samples evaluated across the 17 included studies. Each row corresponds to one sample. The third column (“Exclude”) indicates whether the sample was retained for analysis (“Include”) or removed (“exclude”), with the reason for exclusion noted in parentheses for the 9 excluded samples (e.g., biopsy characterized as prostatic intraepithelial neoplasia, low nFeature, not present in the original paper’s main supplementary table, or irregular UMAP distribution). Of 172 samples collected from the original publications, 163 samples are marked “Include” and 9 are excluded. Additional columns provide: study identifier, patient identifier, sample identifier, internal sample ID, tissue site, histological classification, Gleason score, Gleason grade group, disease state, clinical variables (age, BMI, race, comorbidities), surgical margin status, and sequencing library information.

**Supplementary Table 3. Per-sample quality control summary**. Per-sample QC metrics and cell counts for all 163 included samples. Columns include: study identifier (Study), sample name (Sample), internal sample ID (ID), inclusion status (Exclude), number of cells before QC filtering (Cells_before_QC), median genes and UMIs before QC, percent mitochondrial transcripts before QC, QC filtering thresholds applied, number of cells after QC filtering, median genes and UMIs after QC, number of doublets detected (scDblFinder), doublet rate (%), number of singlets retained, median genes and UMIs in singlets, low-quality cell type annotation flag, Gleason score, Gleason grade group, patient identifier, and number of cells retained after removal of low-quality cell type annotations. QC thresholds applied uniformly: ≥250 detected genes, ≥500 UMI counts, <20% mitochondrial transcripts.

**Supplementary Table 4. Raw scib-metrics benchmark scores — all cell types**. Quantitative benchmarking results from the scib-metrics framework (Luecken et al. 2022, *Nature Methods*) applied to all cell types (2,000 highly variable genes, 50 principal components; batch key: study of origin; label key: cell type annotation). Rows represent six integration methods (Unintegrated, Harmony, Scanorama, BBKNN, scVI, scANVI). Columns represent individual metrics for biological conservation (Isolated labels, KMeans NMI, KMeans ARI, Silhouette label, cLISI) and batch correction (BRAS, iLISI, KBET, Graph connectivity, PCR comparison), plus sub-scores and total score (Total = 0.4 × Batch correction + 0.6 × Bio conservation). All values are raw scores from 0 (worst) to 1 (best). N/A for BBKNN in PCR comparison reflects that BBKNN is a graph-only method and does not produce a corrected embedding.

**Supplementary Table 5. Min-max scaled scib benchmark scores — all cell types**. Same as Supplementary Table 4, with each metric column independently rescaled by min-max normalization (0 = worst-performing method, 1 = best-performing method within each metric). These scaled values correspond to Figure 5c and were used for final method selection. scANVI achieved the highest total score (0.74).

**Supplementary Table 6. Raw scib benchmark scores for luminal cells**. scib-metrics benchmarking results for the luminal cell subset (3,000 highly variable genes, 30 principal components; batch key: study of origin; label key: ETS molecular subgroup). Eight integration methods are evaluated: Unintegrated (PCA baseline), CCA (Seurat v5), RPCA (Seurat v5), Harmony, Scanorama, BBKNN, scVI, and scANVI. Biological conservation metrics: Isolated label score, KMeans NMI, KMeans ARI, Silhouette label score, and cLISI. Batch correction metrics: BRAS, iLISI, kBET, Graph connectivity, and PCR comparison. PCR comparison is not applicable for BBKNN, which does not produce a corrected high-dimensional embedding. Aggregate scores are computed as: Total = 0.4 × Batch correction + 0.6 × Biological conservation. Values are raw (unscaled) scores.

**Supplementary Table 7. Min-max scaled scib benchmark scores for luminal cells**. Same as Supplementary Table 6, with per-column min-max normalization applied across all eight methods (0 = worst-performing method, 1 = best-performing method for each metric). Scaled values correspond to the numbers displayed within circles in Figure 5d. Aggregate scores (Batch correction, Biological conservation, Total) are computed from the scaled per-metric scores. scANVI achieves the highest total aggregate score (0.78).

