## Supplementary figures and images for "A harmonized single-cell RNA-seq atlas of human localized and metastatic prostate cancers and benign tissues"

### Supplemental Figures 1 - 4

#### Cho et al. Supp Figure 1

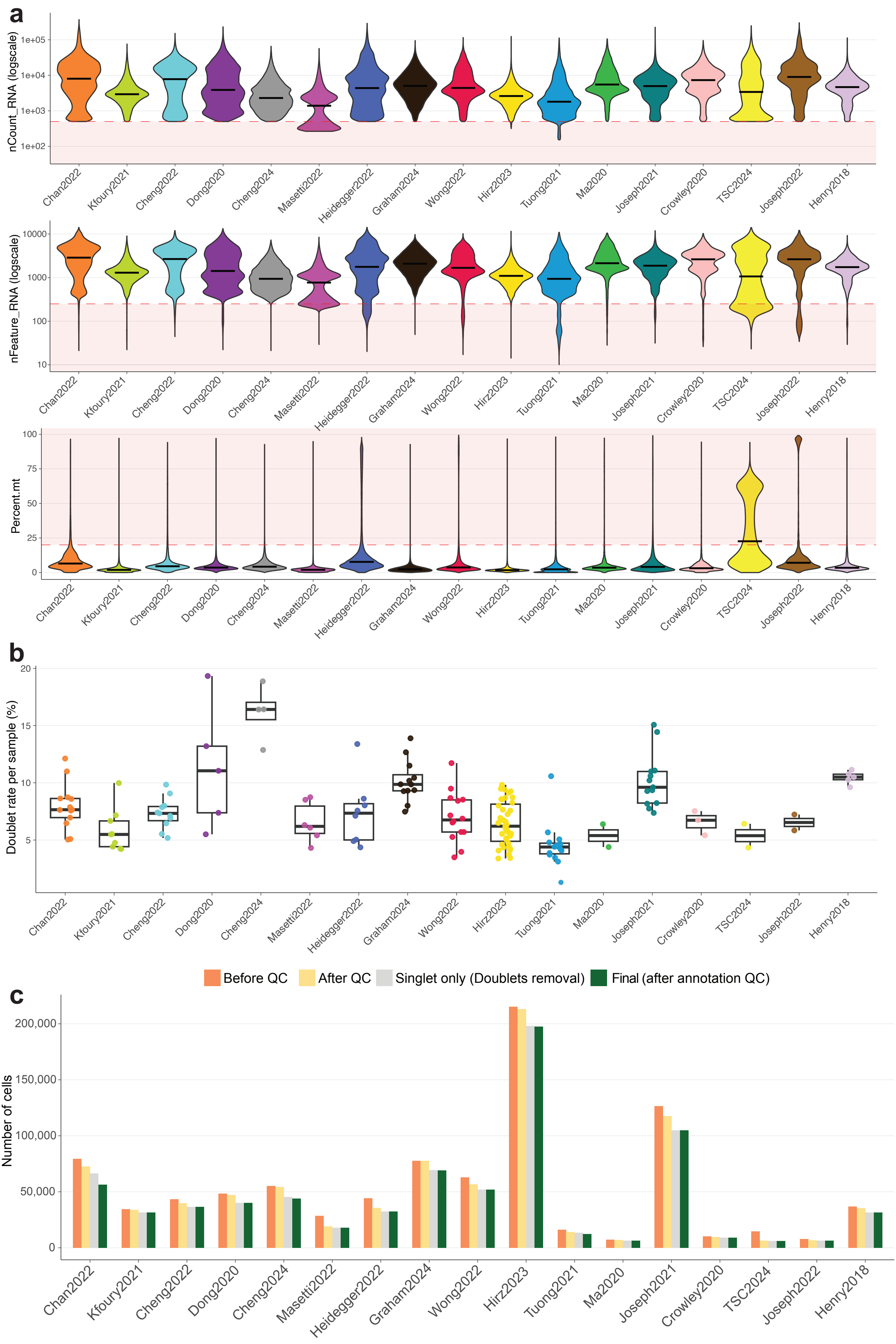

#### Cho et al. Suppl Figure 2

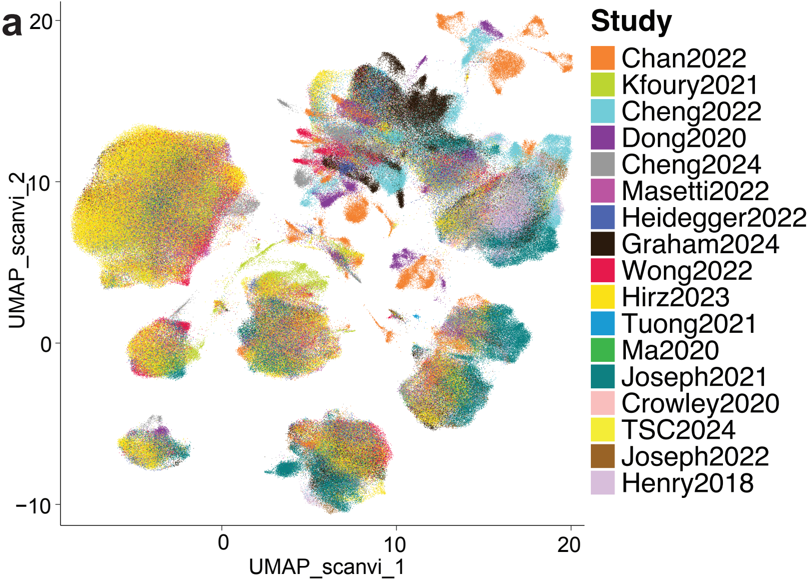


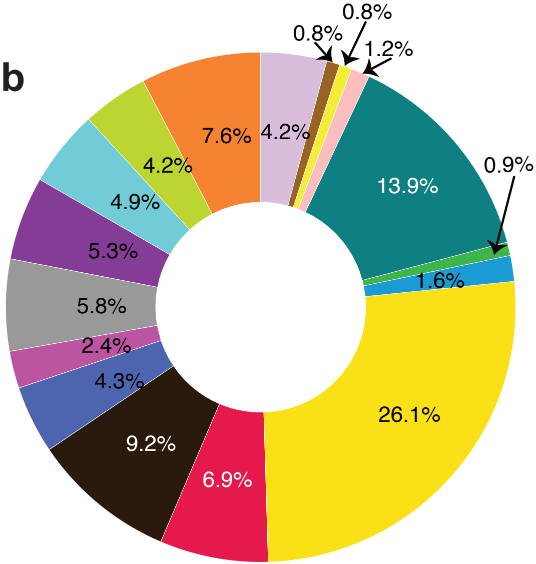

#### Cho et al. Suppl Figure 3

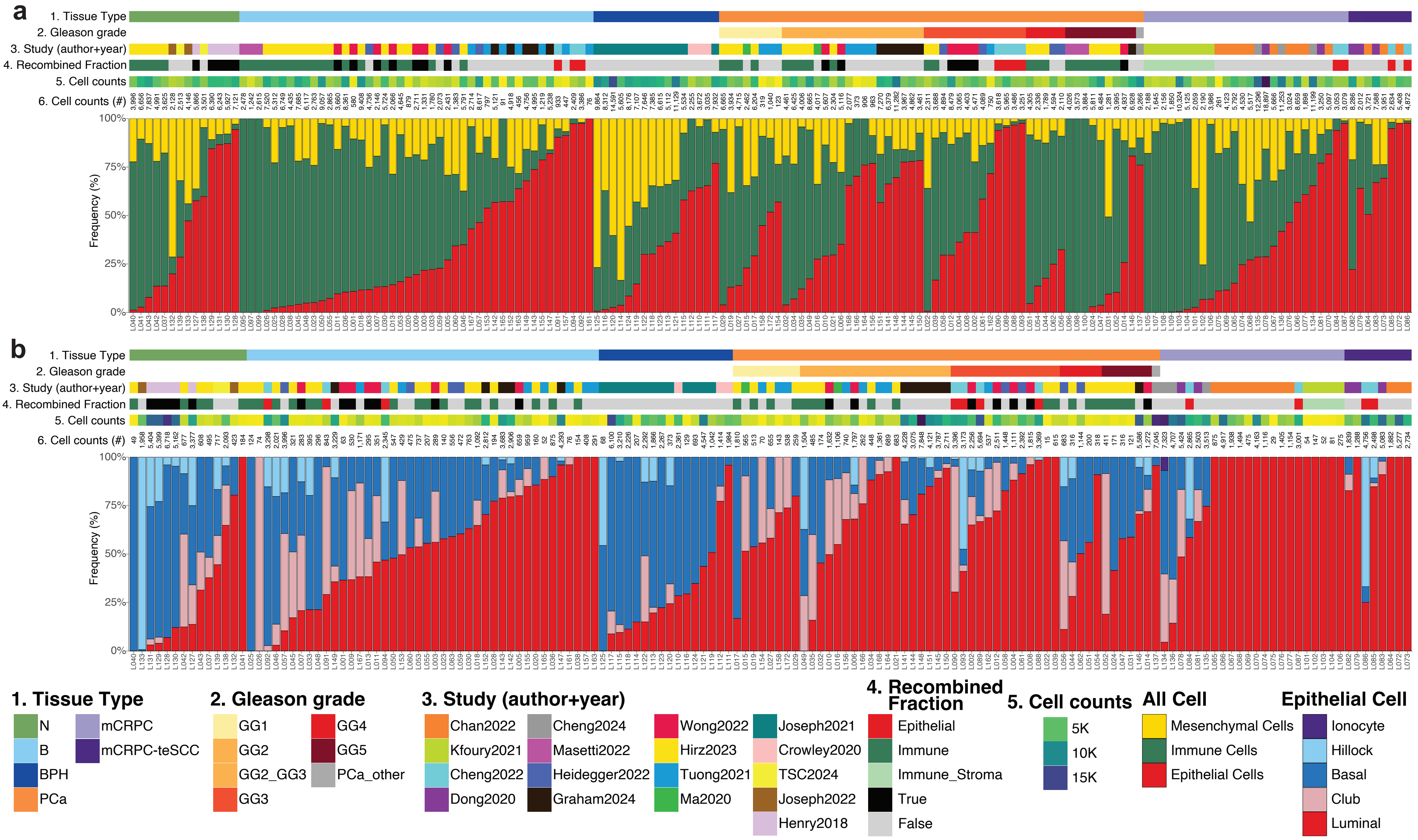

#### Cho et al. Suppl Figure 4

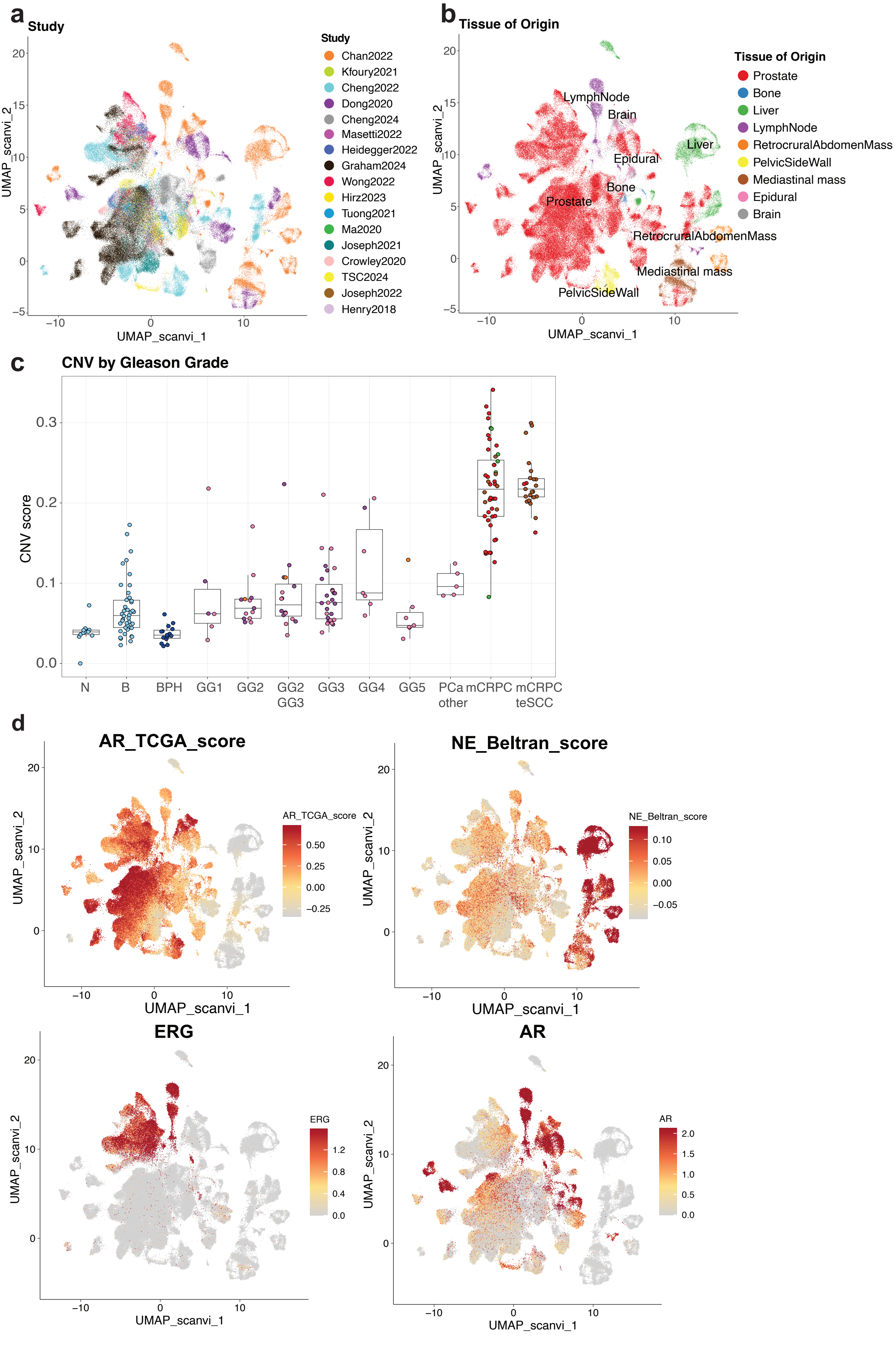
